# Reconstructing imagined letters from early visual cortex reveals tight topographic correspondence between visual mental imagery and perception

**DOI:** 10.1101/277020

**Authors:** Mario Senden, Thomas Emmerling, Rick van Hoof, Martin Frost, Rainer Goebel

## Abstract

Visual mental imagery is the quasi-perceptual experience of “seeing in the mind’s eye”. While a tight correspondence between imagery and perception in terms of subjective experience is well established, their correspondence in terms of neural representations remains insufficiently understood. In the present study, we exploit the high spatial resolution of functional magnetic resonance imaging (fMRI) at 7T, the retinotopic organization of early visual cortex, and machine learning techniques to investigate whether visual imagery of letter shapes preserves the topographic organization of perceived shapes. Sub-millimeter resolution fMRI images were obtained from early visual cortex in six subjects performing visual imagery of four different letter shapes. Predictions of imagery voxel activation patterns based on a population receptive field encoding model and physical letter stimuli provided first evidence in favor of detailed topographic organization. Subsequent visual field reconstructions of imagery data based on the inversion of the encoding model further showed that visual imagery preserves the geometric profile of letter shapes. These results open new avenues for decoding as we show that a denoising autoencoder can be used to pretrain a classifier purely based on perceptual data before fine-tuning it on imagery data. Finally, we show that the autoencoder can project imagery-related voxel activations onto their perceptual counterpart allowing for visually recognizable reconstructions even at the single-trial level. The latter may eventually be utilized for the development of content-based BCI letter-speller systems.

## Introduction

Visual mental imagery refers to the fascinating phenomenon of quasi-perceptual experiences in the absence of external stimulation (Thomas 1999). The capacity to imagine has important cognitive implications and has been linked to working memory, problem solving and creativity (Albers et al. 2013; Kozhevnikov et al. 2013). Yet, the precise degree of similarity to perception remains insufficiently understood. Several functional magnetic resonance imaging (fMRI) studies indicated that imagery activates cortical networks that are also activated during corresponding perceptual tasks (Kosslyn et al. 1997; Goebel et al. 1998; Ishai et al. 2000; O’Craven and Kanwisher 2000; Ganis et al. 2004; Mechelli et al. 2004), lending credence to the notion that imagery resembles perception. Applying multi-voxel pattern analyses (MVPA) furthermore enabled the decoding of feature-specific imagery content related to orientations (Harrison and Tong 2009; Albers et al. 2013), motion (Emmerling et al. 2016), objects (Reddy et al. 2010; Cichy et al. 2012; Lee et al. 2012), shapes (Stokes et al. 2009, 2011), and scenes (Johnson and Johnson 2014).

The MVPA approach has recently been criticized on the grounds that it does not rely on an explicit encoding model of low-level visual features, leaving open the possibility that classification may have resulted from confounding factors such as attention (Naselaris et al. 2015). To overcome this limitation, the authors developed an encoding model based on Gabor wavelets which they fit to voxel activations measured in response to perception of artworks. Subsequently, they used the estimated encoding model to identify an imagined artwork from a set of candidates by comparing voxel activations empirically observed in response to imagery with those predicted from encoding each candidate (Naselaris et al. 2015).

While this study constitutes a major methodological advancement and largely defuses the aforementioned confounds, a complex encoding model allows only for limited inferences regarding the similarity of perception and imagery with respect to any particular feature. It is, for instance, conceivable that the largest contributor to image identification stemmed from an unspecific top-down modulation of salient regions in the imaged artwork with crude retinotopic organization. That is, activations in response to mental imagery might have been co-localized to highly salient regions of the image (without otherwise resembling it) and this might have been sufficient for image identification.

Indeed, results from studies reconstructing the visual field from fMRI data leveraging the retinotopic organization of early visual cortex give the impression that the retinotopic organization of mental imagery is rather diffuse. For instance, while seminal work has been conducted detailing the ability to obtain straightforwardly recognizable reconstructions of perceived physical stimuli (Thirion et al. 2006; Miyawaki et al. 2008; Schoenmakers et al. 2013); similar successes have not been repeated for imagery. Retinotopy-based reconstructions of imagined shapes have so far merely been co-localized with the region of the visual field where they were imagined but bore no visual resemblance to their geometry (Thirion et al. 2006).

However, unless imagery of an object preserves the object’s geometry, it is unlikely it would preserve any of its more fine-grained features. Prior to studying the role such features may play for imagery it is thus pivotal to empirically establish precise topographic correspondence between imagery and perception. Utilizing the high spatial resolution offered by 7T fMRI and the straightforwardly invertible population receptive field model (Dumoulin and Wandell 2008), we provide new evidence that imagery-based reconstructions of letter shapes are recognizable and preserve their physical geometry. This supports the notion of tight topographic correspondence in early visual cortex. Such correspondence opens new avenues for decoding as we show that using a denoising autoencoder it is possible to pretrain a classifier purely based on perceptual data before fine-tuning it on imagery data. Finally, we show that an autoencoder can project imagery-related voxel activations onto their perceptual counterpart allowing for recognizable reconstructions even at a single-trial level. The latter could open new frontiers for brain-computer interfaces (BCIs).

## Materials and Methods

### Participants

Six participants (2 female, age range = (21 -49), mean age = 30.7) with normal or corrected-to-normal visual acuity took part in this study. All participants were experienced in undergoing high field fMRI experiments, gave written informed consent and were paid for participation. All procedures were conducted with approval from the local Ethical Committee of the Faculty of Psychology and Neuroscience at Maastricht University.

### Stimuli and Tasks

Each participant completed three training sessions to practice the controlled imagery of visual letters prior to a single scanning session which comprised four experimental (imagery) runs of ∼11 minutes and one control (perception) run of ∼ 9 minutes as well as one pRF mapping run of ∼16 minutes.

#### Training Session and Task

Training sessions lasted ca. 45 minutes and were scheduled one week prior to scanning. Before the first training session, participants filled in the Vividness of Visual Imagery Questionnaire (VVIQ; Marks, 1973) and the Object-Spatial Imagery and Verbal Questionnaire (Blazhenkova and Kozhevnikov 2009). These questionnaires measure the subjective clearness and vividness of imagined objects and cognitive styles during mental imagery, respectively. In each training trial, participants saw one of four white letters (‘H’, ‘T’, ‘S’, or ‘C’) enclosed in a white square guide box (8° by 8° visual angle) on grey background and a red fixation dot in the center of the screen (see figure 1). With the onset of the visual stimulation, participants heard a pattern of three low tones (note C5) and one high tone (note G5) that lasted 1000 ms. This tone pattern was associated with the visually presented letter with specific patterns randomly assigned for each participant. After 3000 ms the letter started to fade out until it completely disappeared at 5000 ms after trial onset. The fixation dot then turned orange and participants were instructed to maintain a vivid image of the presented letter. After an 18 second imagery period, the fixation dot turned white and probing started. With an inter-probe-interval of 1500 ms (jittered by ±200 ms) three white probe dots appeared within the guide box. These dots were located within the letter shape or outside of the letter shape (however, always within the guide box). Participants were instructed to indicate by button press whether a probe was located inside or outside the imagined letter shape (cf. Podgorny and Shepard 1978). Depending on the response, the fixation dot turned red (incorrect) or green (correct) before turning white again as soon as the next probe was shown. The positions of the probe dots were randomly chosen such that they had a minimum distance of 0.16° and a maximum distance of 0.32° of visual angle from the edges of the letter (and the guide box), both for inside and outside probes. This ensured similar task difficulty across trials. A resting phase of 3000 ms or 6000 ms followed the three probes. At the beginning of a training run all four letters were presented for 3000 ms each, alongside the associated tone pattern (reference phase). During one training run, each participant completed 16 pseudo-randomly presented trials. In each training session, participants completed two training runs during which reference letters were presented in each trial (described above) and two training runs without visual presentation (i.e. the tone pattern was the only cue for a letter). Participants were instructed to maintain central fixation throughout the entire run. After the training session, participants verbally reported the imagery strategies they used.

**Figure 1:**
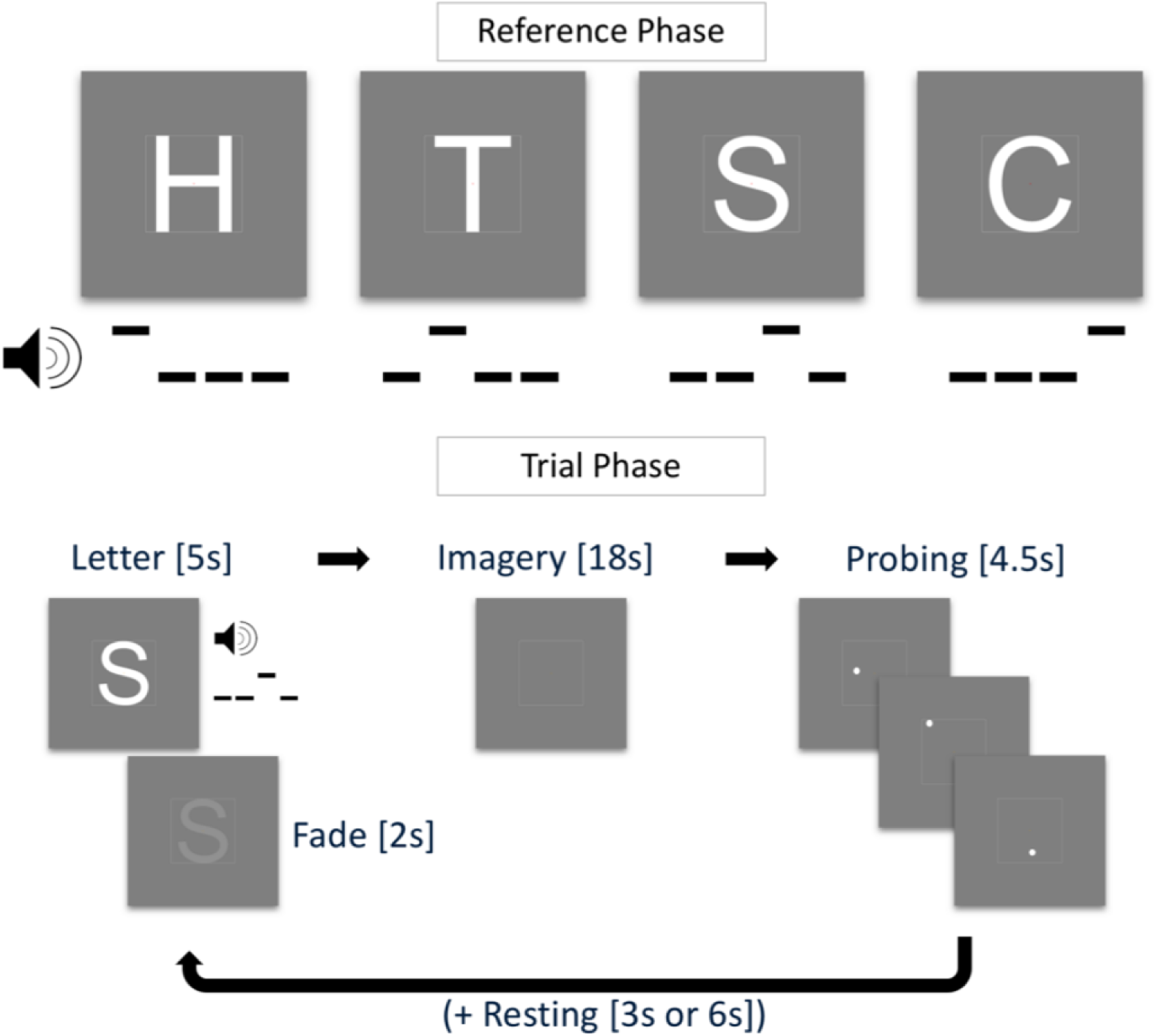
Training task. In the reference phase (top), four letters H, T, ‘S’ & ‘C’ were paired with a tone pattern. In the trial phase (bottom), the tone pattern was played and the letter shown for 5s (fading out after 3s) followed by an imagery period of 18s, a probing period of 4.5s, and a resting period of 3s or 6s

#### Imagery Runs

Imagery runs were similar to the training task with changes to the probing phase and the timing of the trial phase. After the reference phase in the beginning of each run, there was no visual stimulation other than the fixation dot and the guide box. Imagery phases started when participants heard the tone pattern and the fixation dot turned orange. Imagery phases lasted 6s. Participants were instructed to imagine the letter associated with the tone pattern as vividly and accurately as possible. The guide box aided the participant by acting as a reference for the physical dimensions of the letter. The resting phases that followed each imagery phase lasted 9s or 12s. There was no probing phase in normal trials. In each experimental run, there were 32 normal trials and two additional catch trials which entailed a probing phase of four probes. There was no visual feedback for the responses in the probing phase (the fixation dot remained white). Data from the catch trials were not included in the analyses.

#### Perception Run

To measure brain activation patterns in visual areas during the perception of the letters used in the imagery runs we recorded one perception run during the scanning session. The four letters were visually presented using the same trial timing parameters as in the experimental runs. There were neither reference nor probing phases. Letters were presented for the duration of the imagery phase (6s) and their shape was filled with a flickering checkerboard pattern (10 Hz). No tone patterns were played during the perception run. The recorded responses were also used to train a denoising autoencoder (see below).

#### pRF mapping

A bar aperture (1.33° wide) revealing a flickering checkerboard pattern (10 Hz) was presented in four orientations. For each orientation the bar covered the entire screen in 12 discrete steps (each step lasting 3 seconds). Within each orientation the sequence of steps (and hence of the locations) was randomized (cf. Senden et al. 2014). Each orientation was presented six times.

### Stimulus Presentation

The bar stimulus used for pRF mapping was created using the open source stimulus presentation tool BrainStim (http://svengijsen.github.io/BrainStim/).Visual and auditory stimulation in the imagery and perception runs were controlled with PsychoPy (version 1.83.03; Peirce 2007). Visual stimuli were projected on a frosted screen at the top end of the scanner table by means of an LCD projector (Panasonic, No PT-EZ57OEL; Newark, NJ, USA). Auditory stimulation was presented using MR-compatible insert earphones (Sensimetrics, Model S14; Malden, MA, USA). Responses to the probes were recorded with MR-compatible button boxes (Current Designs, 8-button response device, HHSC-2×4-C; Philadelphia, USA).

### Magnetic resonance imaging

We recorded anatomical and functional images with a Siemens Magnetom 7 Tesla scanner (Siemens; Erlangen, Germany) and a 32-channel head-coil (Nova Medical Inc.; Wilmington, MA, USA). Prior to functional scans, we used a T1-weighted magnetization prepared rapid acquisition gradient echo (Marques et al. 2010) sequence [240 sagittal slices, matrix = 320 320, voxel size = 0.7 by 0.7 by 0.7 mm^3^, first inversion time TI1 = 900 ms, second inversion time TI2 = 2750 ms, echo time (TE) = 2.46 ms, repetition time (TR) = 5000 ms, first nominal flip angle = 5°, second nominal flip angle = 3°] to acquire anatomical data. For all functional runs we acquired high-resolution gradient echo (T2* weighted) echo-planar imaging (Moeller et al. 2010) data (TE = 26 ms, TR = 3000 ms, generalized auto-calibrating partially parallel acquisitions (GRAPPA) factor = 3, multi-band factor = 2, nominal flip angle = 55°, number of slices = 82, matrix = 186 by 186, voxel size = 0.8 by 0.8 by 0.8 mm^3^). The field-of-view covered occipital, parietal, and temporal areas. Additionally, before the first functional scan we recorded five functional volumes with opposed phase encoding directions to correct for EPI distortions that occur at higher field strengths (Andersson et al. 2003).

### Processing of (f)MRI data

We analyzed anatomical and functional images using BrainVoyager 20 (version 20.0; Brain Innovation; Maastricht, The Netherlands) and custom code in MATLAB (version 2017a; The Mathworks Inc.; Natick, MA, USA). We interpolated anatomical images to a nominal resolution of 0.8 mm isotropic to match the resolution of functional images. In the anatomical images, the grey/white matter boundary was detected and segmented using the advanced automatic segmentation tools of BrainVoyager 20 which are optimized for high-field MRI data. A region-growing approach analyzed local intensity histograms, corrected topological errors of the segmented grey/white matter border and finally reconstructed meshes of the cortical surfaces (Kriegeskorte and Goebel 2001; Goebel et al. 2006). The functional images were corrected for motion artefacts using the 3D rigid body motion correction algorithm implemented in BrainVoyager 20 and all functional runs were aligned to the first volume of the first functional run. We corrected EPI distortions using a method similar to that described in Andersson, Skare, and Ashburner (2003) and the ‘topup’ tool implemented in FSL (Smith et al. 2004). The pairs of reversed phase encoding images recorded in the beginning of the scanning session were used to estimate the susceptibility-induced off-resonance field and correct the distortions in the remaining functional runs. After this correction, functional data were high-pass filtered using a general linear model (GLM) Fourier basis set of three cycles sine/cosine, respectively. This filtering included a linear trend removal. Finally, functional runs were co-registered and aligned to the anatomical scan using an affine transformation (9 parameters) and z-normalized to eliminate signal offsets and inter-run variance.

### pRF Mapping and region-of-interest definition

For each subject, we fit location and size parameters of an isotropic Gaussian population receptive field model (Dumoulin and Wandell 2008) by performing a grid search. In terms of pRF location, the visual field was split into a circular grid of 100 by 100 points whose density decays exponentially with eccentricity. Receptive field size exhibits a linear relationship with eccentricity with the exact slope depending on the visual area (Freeman and Simoncelli 2011). For this reason, we explored slopes in the range from 0.1 to 1 (step = 0.1) as this effectively allows for exploration of a greater range of receptive field sizes (10 for each unique eccentricity value). We used the pRF mapping tool from the publicly available Computational Neuroimaging Toolbox (https://github.com/MSenden/CNI_toolbox). Polar angle maps resulting from pRF fitting were projected onto inflated cortical surface reconstructions and used to define regions-of-interest (ROIs) for bilateral visual areas V1, V2, and V3. The resulting surface patches from the left and right hemisphere were projected back into volume space (from -1 mm to +3 mm from the segmented grey/white matter boundary). Volume ROIs were then defined for V1, V2, V3, and a combined ROI (V1V2V3).

### Voxel patterns

All our analyses and reconstructions are based on letter-specific spatial activation profiles of voxel co-activations; i.e. voxel patterns. Voxel patterns within each ROI were obtained for both perceptual and imagery runs. First, for each run, single trial letter-specific voxel patterns were obtained by averaging BOLD activations in the range from +2 until +3 volumes following trial onset and z-normalizing the result. This lead to a total of 8 (one per trial) perceptual and 32 (four runs with 8 trials each) imagery voxel patterns per letter. We furthermore computed perceptual and imagery average voxel patterns per letter by averaging over all single trial patterns (and runs in case of imagery) of a letter and z-normalizing the result. Imagery average voxel patterns were used in an encoding analysis and for assessment of reconstruction quality while perceptual average patterns were used for training a denoising autoencoder (Vincent et al. 2008).

### Encoding analysis

To test the hypothesis that spatial activation profiles of visual mental imagery is geometry-preserving, we tested whether voxel activations predicted from the encoding model (one isotropic Gaussian per voxel) and a physical (binary) stimulus corresponding to the imagined letter provides a significantly better fit with measured voxel activations than predictions from the remaining binary letter stimuli. Specifically, for each participant and ROI, we predicted voxel activations for each of the four letters based on pRF estimates and physical letter stimuli.

### Autoencoder

We trained an autoencoder with a single hidden layer to reproduce average perceptual voxel patterns from noise corrupted versions per subject and ROI. Since the values of voxel patterns follow a Gaussian distribution with a mean of zero and unit standard deviation, we opted for zero-mean additive Gaussian noise with a standard deviation *σ* = 12 for input corruption. The hidden layer consisted of ⌊ 0 .1 *N* _*voxels*_ ⌋ units with rectified linear activation functions. Output units activated linearly. Encoding weights (from input to hidden layer) and decoding weights (from hidden to output layer) were shared such that **W** _d_ = **W** _e_ ^*T*^. We used mean squared distances to measure reconstruction loss and implemented the autoencoder in the TensorFlow library (Abadi et al. 2016) for Python (version 2.7, Python Software Foundation, https://www.python.org/). The autoencoder was trained using the Adam optimizer (Kingma and Ba 2014) with a learning rate of 1 × 10 ^-5^ and a batch size of 100 for 2,000 iterations. In addition to the four average perceptual voxel patterns, we also included an equal amount of noise corrupted mean luminance images to additionally force reconstructions to zero, if the input contained no actual signal. No imagery data was used for training the autoencoder.

### Reconstruction

For each subject and ROI, we reconstructed the visual field from average perceptual and imagery voxel patterns. We obtained weights mapping the cortex to the visual field by inverting the mapping from visual field to cortex given by the population receptive fields. Since **W**_*p R F*_, a v-by-p matrix (with v being the number of voxels and p the number of pixels) mapping a 150-by-150 pixel visual field to the cortex (i.e. *p* = 22500 pixels; after vectorizing the visual field) is not invertible, we minimize the error function

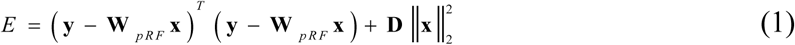

with respect to the input image **x** (a vector of length p). The vector **y** is of length v and reflects a measured voxel pattern. Finally, **D** is a diagonal matrix of the outdegree of each pixel in the visual field which provides pixel-specific scaling of the L2 regularization term 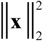 and accounts for cortical magnification. Minimizing equation 1 leads to the expression

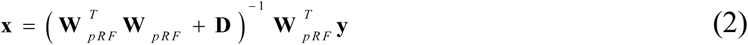

with which we can reconstruct the visual field from voxel patterns. In order to minimize computational cost, we compute the projection matrix 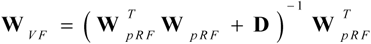 once per ROI and subject rather than performing costly matrix inversion for every reconstruction. For each letter, we assessed the quality of its reconstruction by calculating the correlation between the reconstruction and the corresponding binary letter stimulus. This constitutes a first-level correlation metric. However, since the four letters bear different visual similarities with each other (e.g. ‘S’ and ‘C’ might resemble each other more closely than either resemble ‘H’), we also defined a second-level correlation metric. Specifically, we obtained one vector of all pairwise correlations between physical letter stimuli and a second vector of pairwise correlations between corresponding reconstructions and correlated these two vectors.

### Classification

We replaced the output layer of the pretrained autoencoder with a four-unit (one for each letter) softmax classifier. Weights from the hidden to the classification layer as well as the biases of output units were then trained to classify single trial imagery voxel patterns using cross-entropy as a measure of loss. Note that pretrained weights from input to hidden layer as well as pretrained hidden unit biases remained fixed throughout training of the classifier. These weights and biases were thus dependent purely on perceptual data. This procedure is equivalent to performing multinomial logistic regression on previously established hidden layer representations. Imagery runs were split into training and testing datasets in a leave-one-run-out procedure such that the classifier was repeatedly trained on a total of 96 voxel patterns (8 trials per 4 letters for each of three runs) and tested on the remaining 32 voxel patterns. We again trained the network using the Adam optimizer. However, in this case the learning rate was 1 x 10^−4^, the batch size equal to 96, and training lasted merely 250 iterations.

### Statistical Analysis

Statistical analyses were performed using MATLAB (version 2017a; The Mathworks Inc.; Natick, MA, USA). We used a significance level of *α* = 0 .0 5 (adjusted for multiple comparisons where appropriate) for all statistical analyses.

Behavioral results were analyzed using repeated-measures ANOVA with task (visible or invisible runs) and time as within-subject factors.

For the encoding analysis, we performed a mixed-model regression for the average voxel activations of each imagined letter within each ROI with physical letter as fixed and participant as random factors, respectively. This was followed by a contrast analysis. For each imagined letter the contrast was always between the corresponding physical stimulus and all remaining physical stimuli. For example, when considering voxel activations for the imagined letter ‘H’, a weight of 3 was placed on activations predicted from the physical letter ‘H’ and a weight of −1 was placed on activations predicted from each of the remaining three letters. Since we repeated the analysis for each imagined letter (4) and single region ROI (3), we performed a total of 12 tests and considered results significant at a corrected cutoff of *α* _*c*_ = 0 .0 5 /1 2 = 0 .0 0 4 2 .

In order to evaluate which factors contribute most to first-level reconstruction quality, we performed mixed-model regression with the VVIQ and the OSIVQ spatial and OSIVQ object scores, ROI (using dummy coding, V1 = reference), letter (dummy coding, ‘H’ = reference), and number of selected voxels (grouped by ROI). To assess second-level reconstruction quality, we use the same approach omitting letter as a predictor. Furthermore, we employed a backward elimination stepwise regression procedure to arrive at the most parsimonious set of predictors. That is, we started with the full model and recursively removed non-significant predictors (starting with the largest p-value) until only significant predictors remained in the model.

In order to assess the significance of classification results, we evaluated average classification accuracy across the four runs against a Null-distribution obtained from 1,000 permutations of a leave-one-run out procedure with randomly scrambled labels. We performed this analysis separately for each subject and ROI and consider accuracy results significant if they exceed the 95^th^ percentile of the Null distribution. To statistically evaluate which factors contribute most to classification accuracy, we performed mixed-model regression with the VVIQ and the OSIVQ spatial and OSIVQ object scores, ROI (using dummy coding, V1 = reference), letter (dummy coding, ‘H’ = reference), and number of selected voxels (again grouped by ROI). Finally, As for the analysis of reconstruction quality, we employed a backward elimination stepwise regression procedure to arrive at the most parsimonious set of predictors.

## Results

### Behavioral results

VVIQ and OSIVQ scores for each participant are shown in figure 2. The average score over participants for VVIQ was 4.07 (95% CI [3.71, 4.43]). For the object, spatial, and verbal sub-scales of OSIVQ, average scores were 2.88 (95% CI [2.48, 3.27]), 3.08 (95% CI [2.75, 3.41]), and 3.81 (95% CI [3.33, 4.29]), respectively. Participants reported that they tried to maintain the afterimage of the fading stimulus as a strategy to enforce vivid and accurate letter imagery. Furthermore, participants determined through button presses whether a probe was located inside or outside the letter shape while the letter was either visible or imagined. A repeated-measures ANOVA with task (visible or invisible runs) and time as within-subject factors revealed a statistically significant effect of time on probing accuracy (*F*_*(2,10)*_ = 19.84, *p*≪ 0.001), and no significant difference for task (*F(1,5)* = 1.10, *p* = .341).

**Figure 2:**
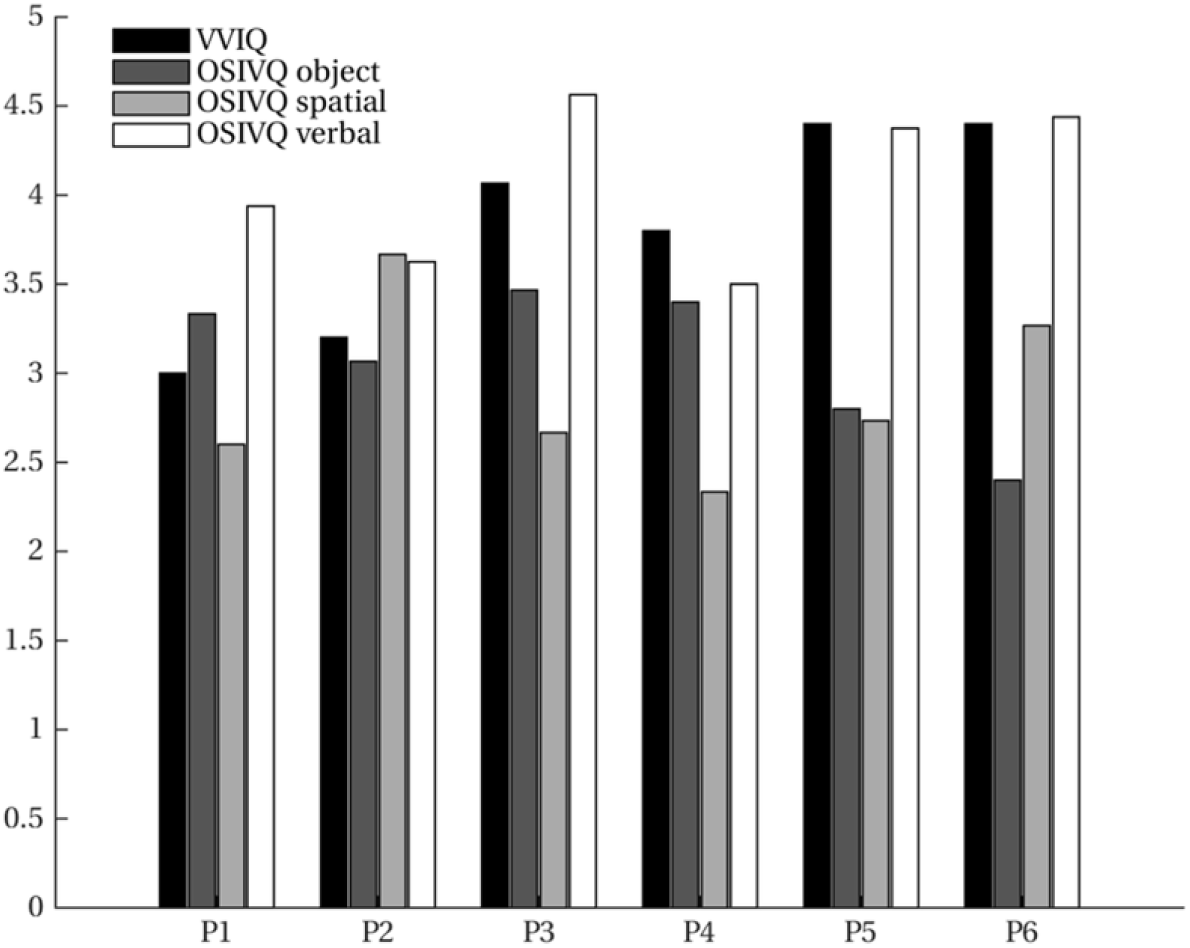
Vividness of visual imagery. Vividness of Visual Imagery Questionnaire (VVIQ) and Object-Spatial Imagery and Verbal Questionnaire (OSIVQ) scores (with the subscales for “object”, “spatial”, and “verbal” imagery styles) are shown for all participants.

### Encoding

For each imagined letter (H, T, S, C) in each single-area ROI (V1, V2, V3), we investigated whether spatial voxel activation profiles can be predicted from a pRF encoding model and the corresponding physical stimulus. That is, for each imagined letter-ROI combination, we ran a mixed-model regression with observed imagery voxel activations (averaged over trials and runs) as outcome variable, predicted voxel activations for each physical letter stimulus as predictors and participants as grouping variable. Since we were specifically interested in testing our hypothesis that the retinotopic organization of imagery voxel activations is sufficiently geometrically specific to distinguish among different imagined letters, we performed contrast analyses between the physical letter corresponding to the imagery and all remaining letters (see Methods for details). Contrasts were significant after applying Bonferroni correction (*α*_*c*_ = 0 .0 0 4 2) for each of the twelve letter-ROI combinations.

Specifically, for V1, predictions based on the physical letter ‘H’ gave a better account of voxel activations observed for the imagery of letter ‘H’ than those based on every other physical letter (*t*_*(2)*_ *=* 32.11, *p =* 0.0004). Similarly for ‘T’ (*t*_*(2)*_ *=* 48.00, *p =* 0.0002), ‘S’ (*t*_*(2)*_ *=* 14.10, *p =* 0.0025), and ‘C’ (*t*_*(2)*_ *=* 29.84, *p =* 0.0006). In the same vain for V2, ‘H’ (*t*_*(2)*_ *=* 25.21, *p =* 0.0008), ‘T’ (*t*_*(2)*_ *=* 67.63, *p =* 0.0001), ‘S’ (*t*_*(2)*_ *=* 19.64, *p =* 0.0013), and ‘C’ (*t*_*(2)*_ *=* 47.48, *p =* 0.0002) and V3, ‘H’ (*t*_*(2)*_ *=* 47.90, *p =* 0.0006), ‘T’ (*t*_*(2)*_ *=* 27.60, *p =* 0.0007), ‘S’ (*t*_*(2)*_ *=* 11.48, *p =* 0.0038), and ‘C’ (*t*_*(2)*_ *=* 32.83, *p =* 0.0005). Figure 3 visualizes these results in the form of boxplots of first-level beta values (i.e. distribution over participants per physical letter) in each letter-ROI combination.

**Figure 3:**
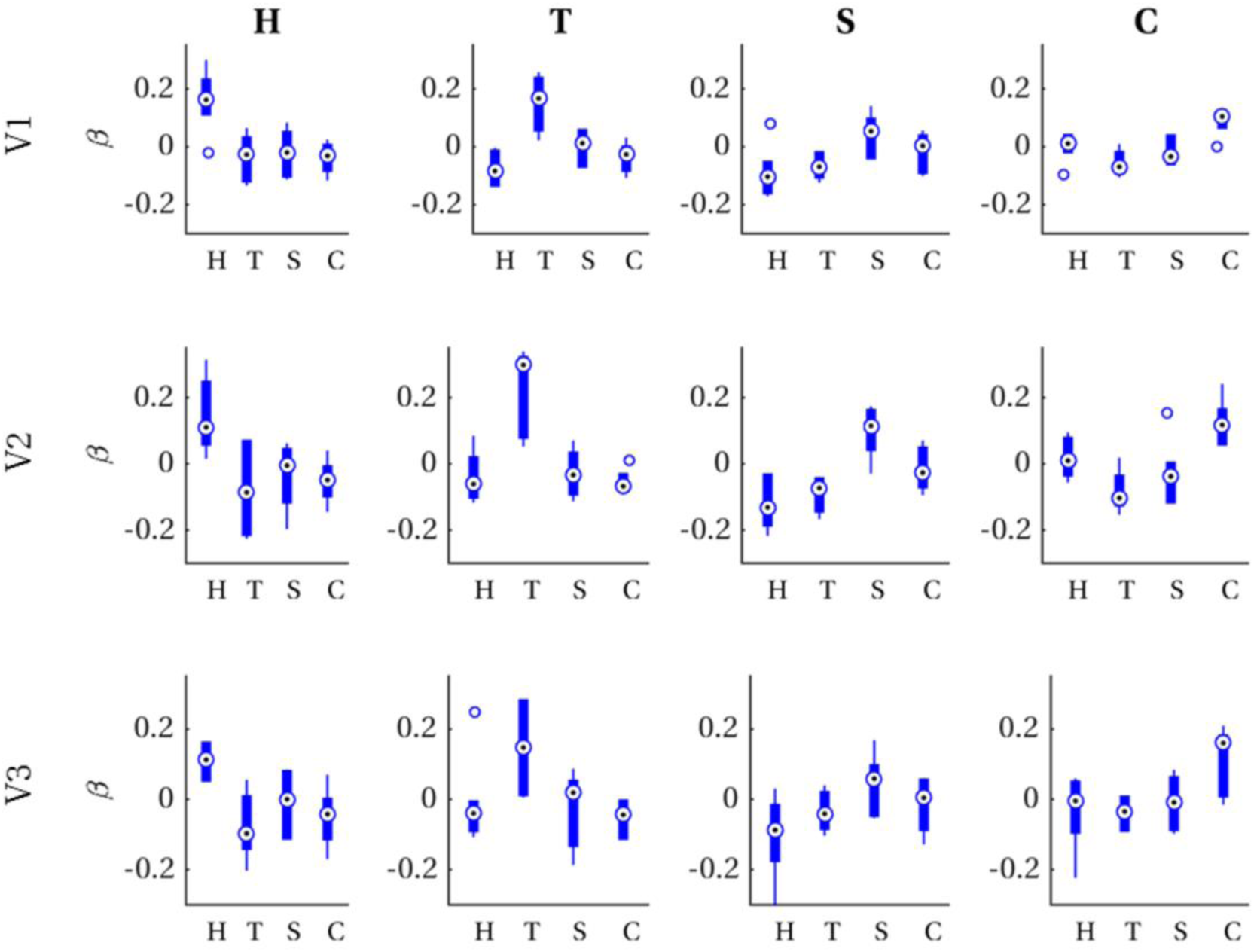
First-level beta distributions. Distribution of first-level beta values (across participants) for voxel patterns predicted from each physical letter (x-axis) for all combinations of ROI (rows) and imagined letters (columns).

### Reconstruction

#### Raw imagery data

We reconstructed the visual field from average imagery voxel patterns in response to each letter (see figures 4 and 5). Mean correlations between reconstructed imagery and physical letters are presented in table 2 (for comparison, table 3 shows correlations between reconstructed perception and physical letters). As can be appreciated from these results as well as the figures, first-level reconstruction quality varies across ROIs as well as across subjects. Differences between subjects might be due to differences in their ability to imagine shapes accurately and vividly as measured by the VVIQ and OSIVQ questionnaires.

**Figure 4:**
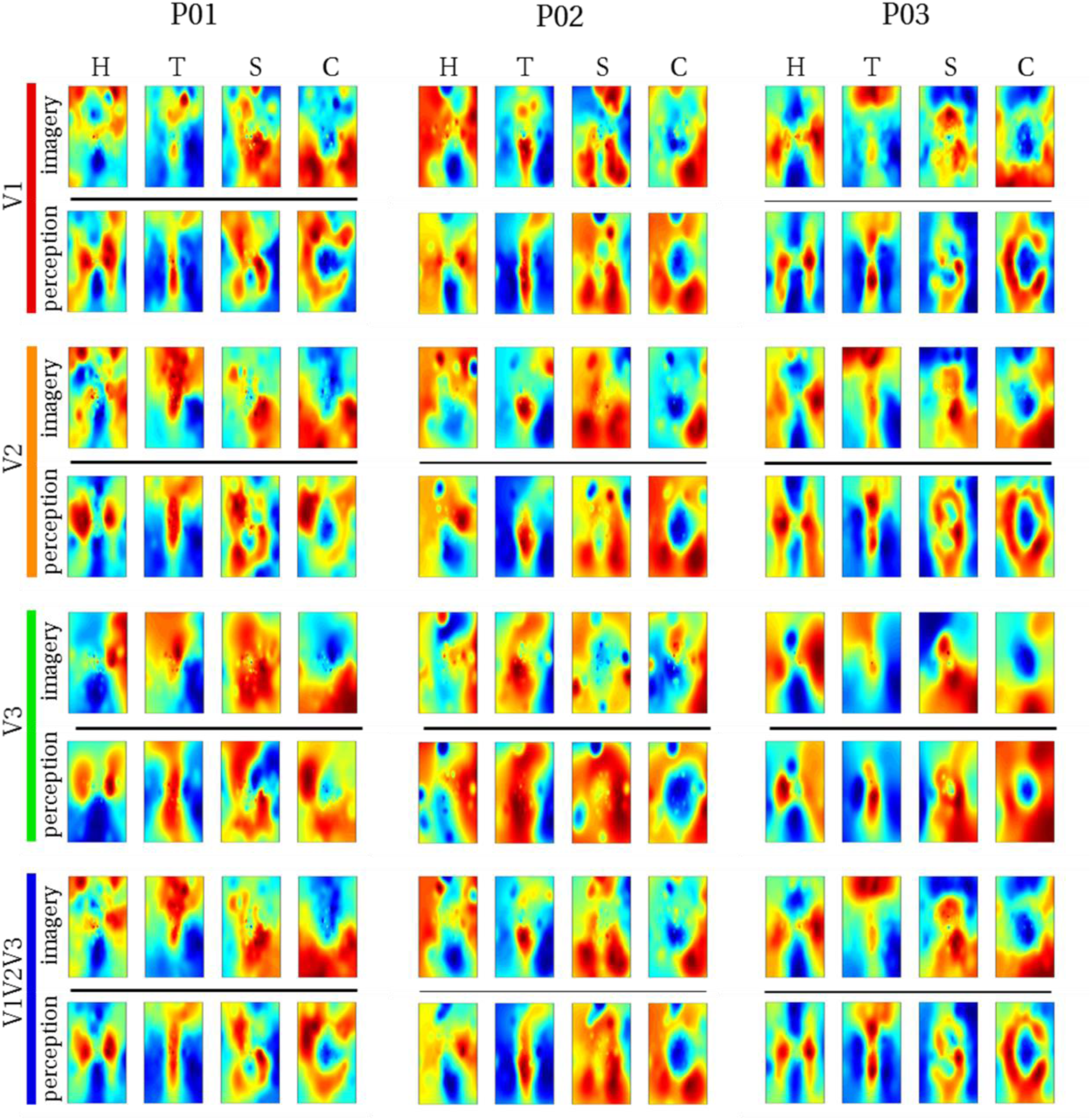
Reconstructed visual field images (participants 1-3). Reconstructed average visual field images are visualized for each ROI of participants one, two, and three. Reconstructions of the remaining three subjects are shown in figure 5. Perceptual as well as imagery voxel patterns were obtained from raw BOLD time-series.

**Figure 5:**
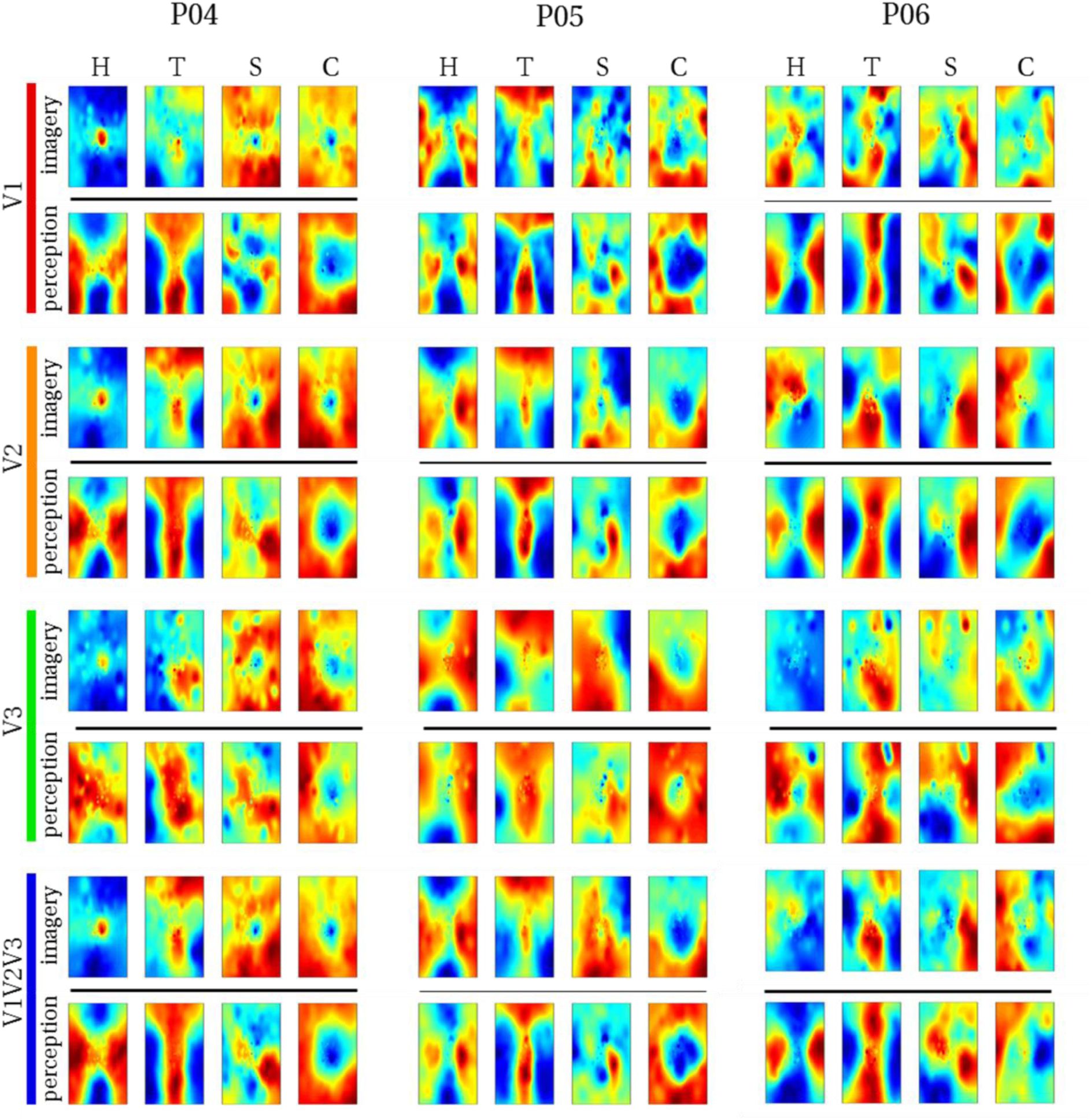
Reconstructed visual field images (participants 4-6). Reconstructed average visual field images are visualized for each ROI of participants four, five, and six. Reconstructions of the remaining three subjects are shown in figure 4. Perceptual as well as imagery voxel patterns were obtained from raw BOLD time-series.

**Table 1.**
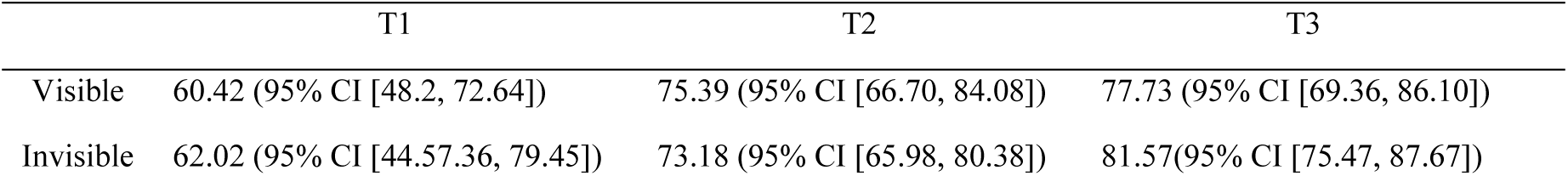
Probing accuracy (averages over participants and time).

**Table 2.** First order correlations between reconstructed imagined letters and physical stimuli (averages over participants).

|  | H | T | S | C |
| --- | --- | --- | --- | --- |
| V1 | 0.24 (95% CI [0.09, 0.40]) | 0.49 (95% CI [0.38, 0.58]) | 0.08 (95% CI [0.03, 0.19]) | 0.14 (95% CI [0.07, 0.21]) |
| V2 | 0.21 (95% CI [0.13, 0.30]) | 0.46 (95% CI [0.36, 0.55]) | 0.10 (95% CI [0.04, 0.17]) | 0.12 (95% CI [0.07, 0.18]) |
| V3 | 0.20 (95% CI [0.10, 0.30]) | 0.28 (95% CI [0.14, 0.46]) | 0.04 (95% CI [0.03, 0.12]) | 0.15 (95% CI [0.01, 0.29]) |
| V1V2V3 | 0.27 (95% CI [0.16, 0.37]) | 0.51 (95% CI [0.45, 0.56]) | 0.12 (95% CI [0.02, 0.21]) | 0.14 (95% CI [0.08, 0.20]) |

**Table 3.** First order correlations between reconstructed perceived letters and physical stimuli (averages over participants).

|  | H | T | S | C |
| --- | --- | --- | --- | --- |
| V1 | 0.40 (95% CI [0.35, 0.44]) | 0.65 (95% CI [0.60, 0.69]) | 0.27 (95% CI [0.15, 0.38]) | 0.32 (95% CI [0.22, 0.40]) |
| V2 | 0.37 (95% CI [0.31, 0.42]) | 0.58 (95% CI [0.50, 0.64]) | 0.19 (95% CI [0.09, 0.29]) | 0.31 (95% CI [0.23, 0.38]) |
| V3 | 0.25 (95% CI [0.19, 0.31]) | 0.41 (95% CI [0.30, 0.51]) | 0.06 (95% CI [-0.06, 0.18]) | 0.27 (95% CI [0.25, 0.30]) |
| V1V2V3 | 0.41 (95% CI [0.36, 0.46]) | 0.63 (95% CI [0.56, 0.68]) | 0.22 (95% CI [0.12, 0.32]) | 0.31 (95% CI [0.24, 0.38]) |

Differences between ROIs might be due to differences with respect to their retinotopy (most likely receptive field sizes) or due to different numbers of voxels included for analysis of each ROI. Only the former would be a true ROI effect. We investigate which factors account for observed correlations (transformed to Fisher z-scores for analyses) by performing a mixed-model regression with questionnaire scores, ROI (using dummy coding, V1 = reference), letter (dummy coding, ‘H’ = reference), and number of selected voxels as predictors. Number of voxels were grouped by ROI. We further performed stepwise model reduction to arrive at the most parsimonious account of our results. In the full model the VVIQ and the OSIVQ spatial and object scores were included but the OSIVQ verbal scores were not since those correlated highly with VVIQ scores (leading to collinearity) and are arguably the least relevant for mental imagery of visual shapes. To further prevent collinearity, we also only included single-area ROIs in this analysis and not the combined ROI. The final model retained number of voxels (*t*_*(64)*_ *=* 3.70, *p* < 0.001) and the OSIVQ object score *t*_*(64)*_ *=* 3.52, *p* < 0.001) as significant quantitative predictors. Furthermore, letter was retained as a significant categorical predictor. Specifically, letter ‘T’ (*t*_*(64)*_ *=* 5.46, *p* ≪ 0.001) presented with significantly improved correlation values over the reference letter ‘H’ whereas letters ‘S’ (*t*_*(64)*_ *=* -3.79, *p* < 0.001) and ‘C’ (*t*_*(64)*_ *=* -2.20, *p* = 0.032) presented with significantly decreased correlation values with respect to the reference.

Next, we examined the second-level correlation metric of reconstruction quality. Correlations between physical and reconstruction pairwise first-level correlation vectors were 0.60 (95% CI [0.28, 0.80], *p =* 0.103) for V1, 0.65 (95% CI [0.34, 0.83], *p =* 0.082) for V2, 0.48 (95% CI [0.15, 0.71], *p =* 0.167) for V3, and 0.64 (95% CI [0.34, 0.83], *p =* 0.084) for V1V2V3, respectively. Finally, we performed a mixed regression to assess which factors account for the observed correlations (again transformed to Fisher z-scores). We included VVIQ and the OSIVQ spatial and object scores, ROI (dummy coding, V1 = reference), and number of selected voxels (grouped by ROI) as predictors and performed stepwise model reduction. The final model retained number of voxels (*t*_*(15)*_ *=* 4.17, *p* < 0.001) and the OSIVQ object score (*t*_*(15)*_ *=* 4.02, *p* = 0.001) as significant predictors.

#### Processed imagery data

Our results confirm that visual mental imagery preserves perceptual topographic organization. This can be leveraged to obtain improved reconstructions of mental imagery. Specifically, an autoencoder trained to retrieve perceptual voxel patterns from their noise corrupted version can be utilized to enhance imagery data. Figures 6 shows how the autoencoder affects first-level reconstruction quality on a single trial basis for V1. As shown in the figure, reconstruction quality was best for ‘T’, followed by ‘H’, ‘C’ and ‘S’. A subject effect is also clearly apparent with participants three and five generally displaying the best results. Finally, imagery reconstruction quality was generally inferior to perception prior to using the autoencoder. However, using the autoencoder pushed imagery reconstruction quality towards perception levels. Indeed, the autoencoder maps imagery voxel patterns onto the corresponding perception voxel patterns it has learned previously. This explains two import observations. First, for some trials, using the autoencoder decreased resemblance to the physical letter. This is especially apparent for participants four and six whose reconstructions were generally not particularly good. Such decrements in reconstruction quality result from imagery voxel patterns in response to one letter falling within the attraction domain of another letter (resembling the activation pattern of that letter slightly more) and hence get mapped onto the wrong pattern. Second, even the few imagery trials whose reconstructions match the physical letter better than the perceptual data the autoencoder was trained were mapped onto the perceptual pattern. A notable example are two trials for the letter ‘S’ by participant two.

**Figure 6:**
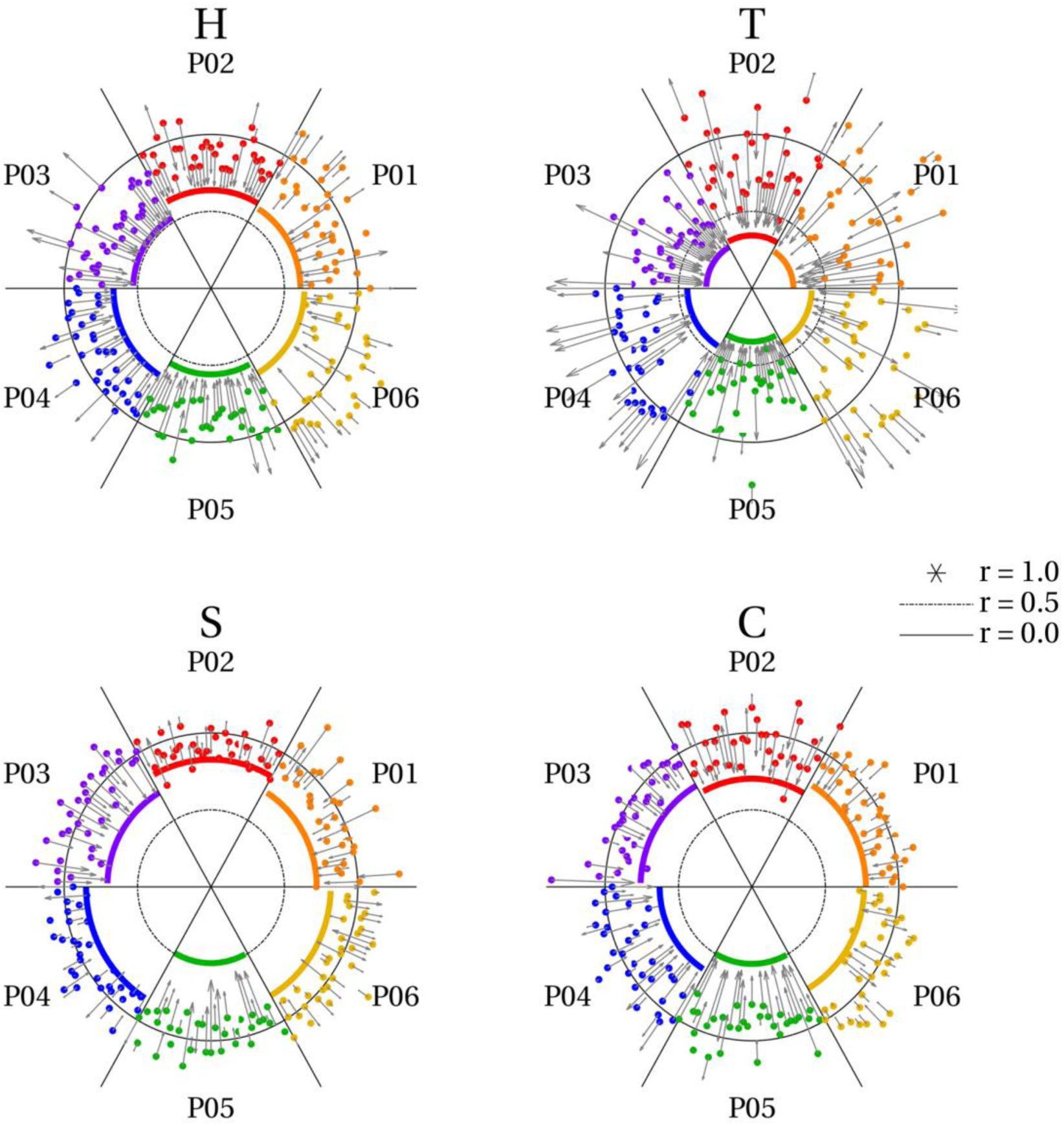
Effect of autoencoder on trial-specific reconstruction quality. The radius of the circles represents reconstruction quality (correlations) with r=1 at the center, r = 0.5 at the inner ring (dash-dot) and r = 0 at the outer ring (solid). Each angle represents an imagery trial with 32 trials per letter and participant. Participants are color coded. Solid colored lines reflect reconstruction quality based on average perceptual voxel patterns of one participant. This constitutes a baseline against which to compare imagery reconstruction quality. Colored dots reflect imagery reconstruction quality for each individual trial of a participant. Finally, arrows reflect the displacement of each of these dots after feeding imagery data through the autoencoder. That is, the tip of the head reflects the new position of the dot after applying the autoencoder. Most points were projected onto the perception-level correlation value and hence approached the center. However, some moved further away from the center.

This implies that reconstruction quality of the perceptual data used to train the autoencoder constitutes an upper limit for imagery when using the autoencoder.

As a general effect, the autoencoder maps imagery voxel patterns onto their perceptual counterpart for most individual trials. Hence, reconstructions of average imagery voxel patterns as well as of individual trials more strongly resemble the corresponding physical letter. Figure 7 shows reconstructions from average imagery voxel patterns after feeding the data through the autoencoder. Figures 8 and 9 show reconstructions of individual trials in a single run of participants three and five, respectively.

**Figure 7:**
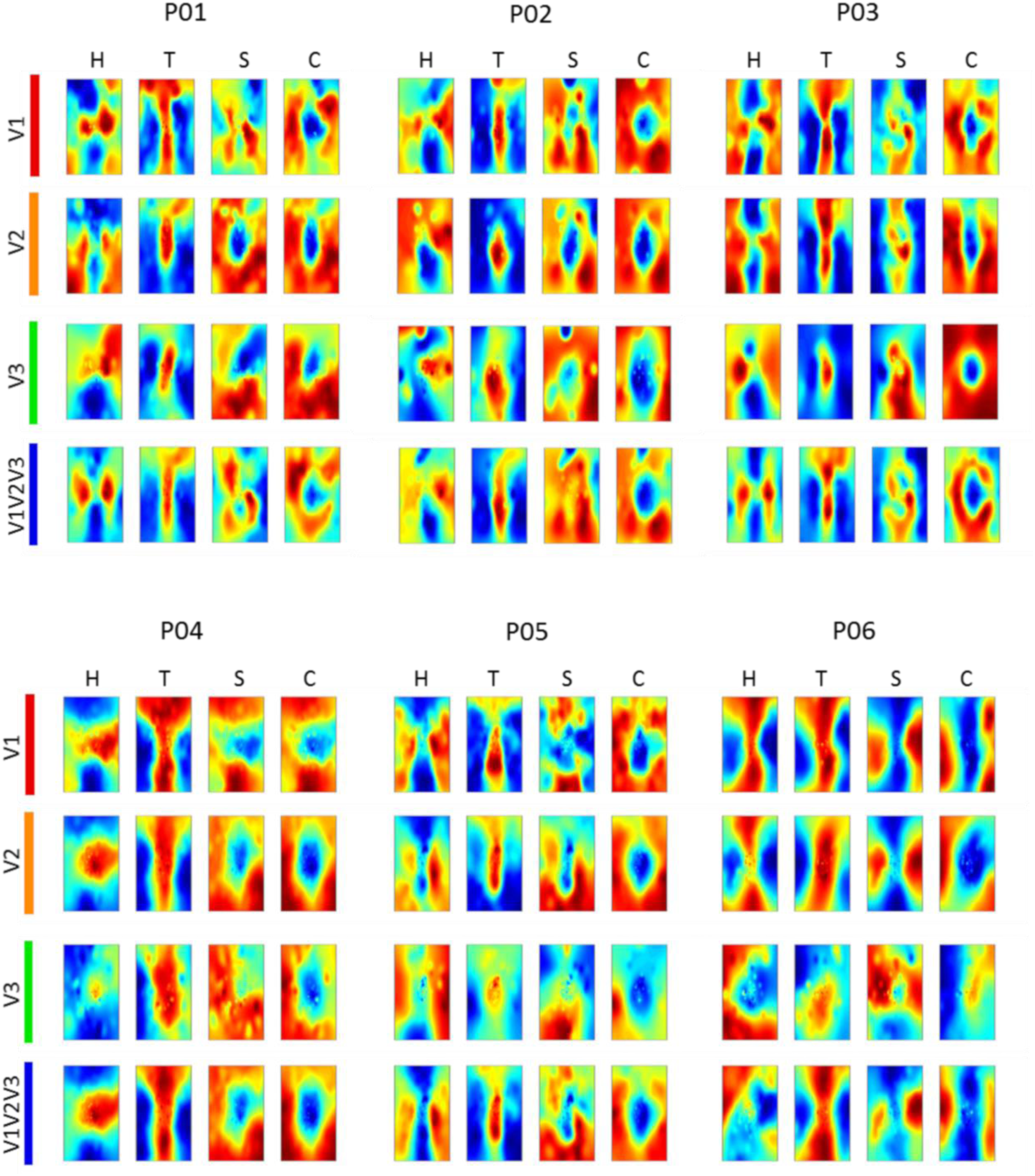
Reconstructed imagery. Reconstructed average visual field images of mental imagery are visualized for each ROI of each participant. Imagery voxel patterns were obtained from cleaned BOLD time-series after feeding raw data through the autoencoder.

**Figure 8:**
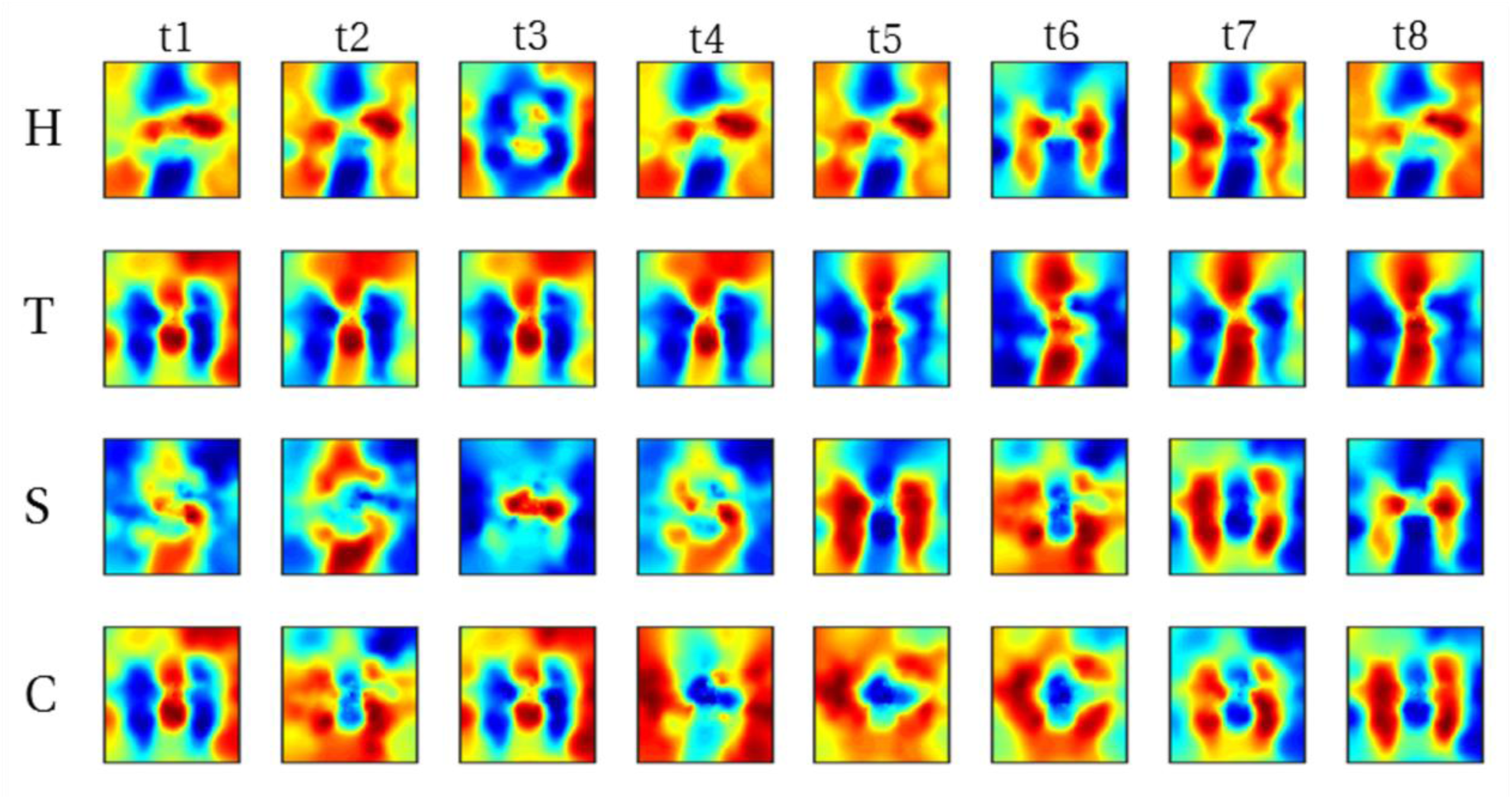
Reconstructed visual field images from denoised single trials in a single run of participant 3. Each run comprised of 8 trials (columns) per letter (rows). Recognizable reconstructions can be obtained for a number (though not all) individual trials.

**Figure 9:**
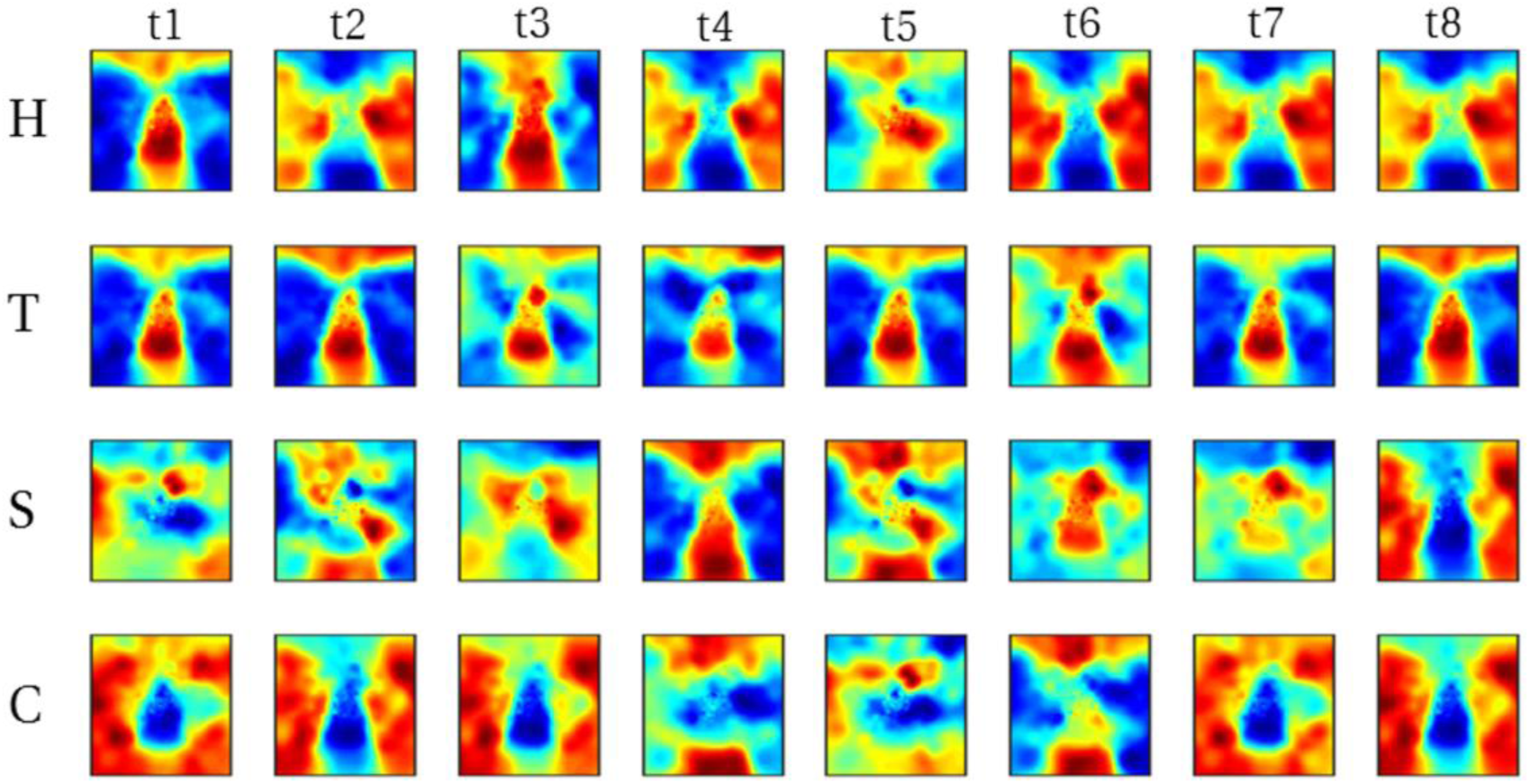
Reconstructed visual field images from denoised single trials in a single run of participant 5. Each run comprised of 8 trials (columns) per letter (rows). Recognizable reconstructions can be obtained for a number (though not all) individual trials.

Obviously, these participants are not representative of the population at large but provide an indication of what is possible for people with a strong ability to imagine visual shapes. If data of participant three were fed through the autoencoder, mean correlation values across trials were 0.39 (95% CI [0.32, 0.45]) for ‘H’, 0.55 (95% CI [0.46, 0.62]) for ‘T’, 0.10 (95% CI [0.04, 0.16]) for ‘S’, and 0.09 (95% CI [0.06, 0.12]) for ‘C’, respectively. For comparison, without using the autoencoder mean correlations were 0.19 (95% CI [0.15, 0.21]) for ‘H’, 0.33 (95% CI [0.28, 0.38]) for ‘T’, -0.02 (95% CI [-0.06, 0.02]) for ‘S’, and 0.02 (95% CI [-0.02, 0.06]) for ‘C’, respectively. If data of participant five were fed through the autoencoder,, mean correlations were 0.28 (95% CI [0.20, 0.35]) for ‘H’, 0.53 (95% CI [0.43, 0.61]) for ‘T’, 0.08 (95% CI [0.00, 0.17]) for ‘S’, and 0.21 (95% CI [0.12, 0.31]) for ‘C’, respectively. For raw imagery data, mean correlations were 0.12 (95% CI [0.08, 0.15]) for ‘H’, 0.32 (95% CI [0.25, 0.37]) for ‘T’, 0.02 (95% CI [-0.01, 0.06]) for ‘S’, and 0.07 (95% CI [0.03, 0.10]) for ‘C’, respectively.

### Classification

Having established support for the hypothesis that activity in early visual cortex in response to imagery exhibits a similar topographical profile as perception, we proceeded to test whether it is possible to pretrain latent representations for an imagery classifier using purely perceptual data. The classifier consists of three layers with the output layer being a softmax classifier stacked onto the hidden layer of an autoencoder pretrained to denoise perceptual voxel patterns (see Methods for details). We trained the classifier on imagery data using a leave-one-run-out procedure; that is, we trained the classifier on three of the four imagery runs and tested classification accuracy on the left-out run. Figure 10 shows average classification accuracies per subject and ROI (including the combined ROI ‘V1V2V3’). For five of the six participants, average classification accuracies exceeded theoretical chance levels (25% correct) as well as the 95^th^ percentile of 1,000 permutation runs (randomly scrambled labels) in all ROIs. For participant six, theoretical chance levels as well as the 95^th^ percentile were (barely) exceeded for V2 only.

**Figure 10:**
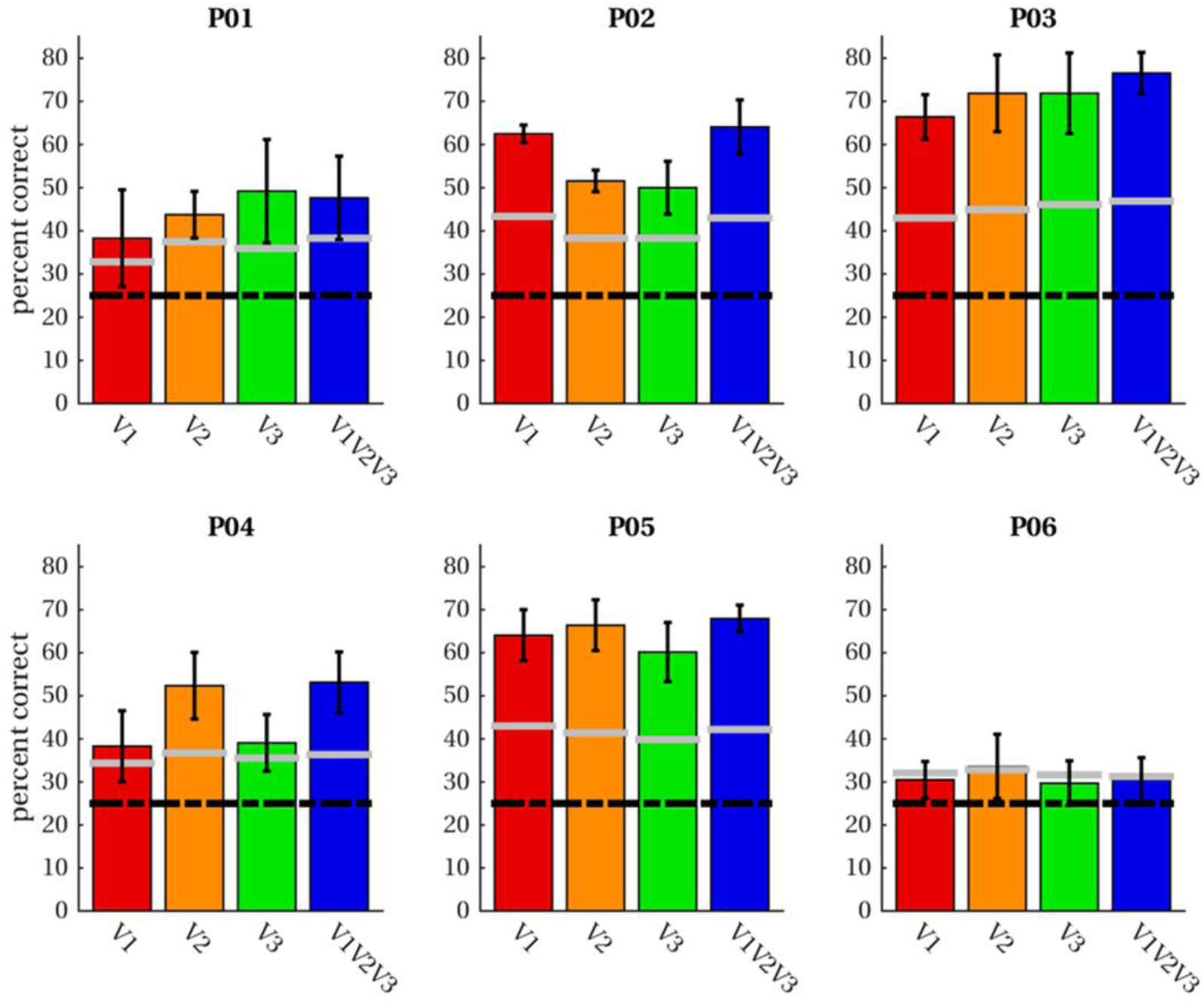
Classification accuracies. Average classification accuracies across four leave-one-out runs of imagery data are given for four ROIs in each participant. Classification was performed for letter-specific voxel patterns averaged in the range from +2 until +3 volumes after trial onset. The black dashed line indicates accuracies expected by chance; grey lines demarcate the 95^th^ percentile of permutation classification accuracies.

We performed a mixed-model regression with the VVIQ and the OSIVQ spatial and OSIVQ object scores, ROI (using dummy coding, V1 = reference), and number of selected voxels (again grouped by ROI) as predictors to assess which factors account for the observed accuracies. We again performed stepwise model reduction to arrive at the most parsimonious account of our results. The final model retains three significant predictors of average classification accuracy: number of voxels (*t*_*(14)*_ *=* 5.37, *p* ≪ 0.001), the object sub-score of OSIVQ (*t*_*(14)*_ *=* 4.57, *p <* 0.001), and the spatial sub-score of OSIVQ (*t*_*(14)*_ *=* 2.95, *p =* 0.011).

## Discussion

The aim of the present study was to investigate whether visual imagery exhibits sufficient topographic organization to preserve the geometry of internally visualized objects. To that end we trained participants to maintain a vivid mental image of four letter shapes. Subsequently, we obtained submillimeter resolution 7T fMRI measurements from early visual cortex while participants viewed or imagined the same letter shapes. Finally, we conducted a series of encoding, reconstruction, and decoding analyses to establish the degree of similarity between imagined and perceived shapes. Our results reveal that an object’s geometry is preserved during visual mental imagery.

Over training sessions all participants reached a high probing accuracy for both imagery and perception trials, showing that they could reliably indicate the location of the invisible letter shape in visual space. The ability to imagine the borders of the letter in absence of visual stimulation suggests participants were able to internally visualize the instructed letter. Next, we showed that patterns of voxel activations predicted by a pRF encoding model and a physical (binary) letter stimulus can account for observed activation patterns in response to mental imagery of the letter corresponding to the physical stimulus. Given that pRF mapping has been shown to accurately predict fMRI responses to visual stimuli (Wandell and Winawer 2015), our results suggest that intrinsic geometric organization of visual experiences are also maintained during visual mental imagery. Our encoding analysis is somewhat reminiscent of that employed by Naselaris et al. (2015) who used a more sophisticated encoding model to identify an imagined artwork from a set of candidates. Given that our encoding model is far simpler, our approach is much more restricted than that detailed in (Naselaris et al. 2015). However, a more restricted approach has the advantage of affording tighter control over the experiment. By focusing on a single feature (retinotopic organization), using stimuli differing solely with respect to their geometric properties, and directly comparing the predictions based on each stimulus regarding the activation profiles in early visual cortex, allowed us to draw specific conclusions regarding the topographic organization of mental imagery.

With respect to reconstructions, we found significant overlap between reconstructed imagery and the physical stimulus in terms of object geometry. While we anticipated this given findings that visual mental imagery exhibits retinotopic organization in early visual cortex (Slotnick et al. 2005; Albers et al. 2013; Pearson et al. 2015), these results were nonetheless exciting because previous reconstructions of mental imagery based on retinotopy did not preserve object geometry (Thirion et al. 2006). Our first-level correlation metric of reconstruction quality revealed that reconstruction quality of letter ‘S’ was significantly reduced while that of letter ‘T’ was significantly improved with respect to that of letter ‘H’. This fits with the notion that stimuli exhibiting finer (coarser) spatial layouts would be harder (easier) to reconstruct. Furthermore, the OSIVQ object score was a significant predictor of first-level reconstruction quality indicating that participants relying on an object-based imagery strategy were generally more successful at imagery of the letter shapes. This finding is also in line with recent observations that neural overlap between imagery and perception in the visual system depends on experienced imagery vividness (Dijkstra et al. 2017).

Interestingly, while inspection of figures 4 and 5 would suggest that reconstruction quality is ROI-specific, ROIs do not constitute a significant predictor of first-level reconstruction quality. Rather, the number of voxels included for any given ROI determined the quality. However, this does not imply that uncritically adding more voxels will definitely lead to higher classification accuracies. We included only those voxels for which pRF mapping yielded a high fit. It is likely that reconstructions benefit from a large number of voxels whose pRF can be estimated to a high degree of precision (i.e. which show a strong spatially selective visual response) rather than a large number of voxels per se. The OSIVQ object score and number of voxels were also significant predictors of second-level reconstruction quality for reasons similar to those just mentioned.

Both our encoding and reconstruction results show that it is possible to extract similar information from perceived and imagined shapes and are strongly indicative of the pictorial nature of vividly experienced mental images (Brogaard and Gatzia 2017). We further show that the tight topographic correspondence between imagery and perception in early visual cortex allows for improved reconstruction and opens new avenues for decoding. Specifically, our results indicate that training a denoising autoencoder on perceptual data creates an attractor landscape with one attractor per perceived letter. Importantly, the resemblance of imagery activation profiles in early visual cortex is sufficiently similar to its perceptual pendant to ensure that activation patterns of a large proportion of imagery trials fall within the attraction domain of the correct letter. The autoencoder then projects these imagery activation patterns onto the corresponding perceptual activation patterns. Though this is not the case for every trial with some being projected onto the wrong perceptual activation pattern pointing to intra-individual fluctuations regarding successful imagery (cf. Dijkstra et al. 2017).

Nevertheless, these projections allow for perception-level reconstruction quality even for individual imagery trials for those participants with good imagery ability. It may be an interesting avenue for future research to study the attractor landscape formed through training the autoencoder and investigate under which conditions imagery trials fall inside or outside the attraction domain of each letter. Our current observations regarding the autoencoder imply that perception provides an upper limit on the achievable reconstruction quality. That is, any improvements of perceptual reconstructions, for instance obtaining a more accurate encoding model by correcting for eye movements during pRF mapping (Hummer et al. 2016), should improve imagery reconstructions as well.

Furthermore, the autoencoder can be utilized to pretrain a classifier purely based on perceptual data before fine-tuning it on imagery data. We showed the feasibility of this approach by using it for classifying imagined letters with a high degree of accuracy from at least one region of interest (between 50% and 70% correct) in five out of six participants. Statistical analyses revealed that both the OSIVQ object and OSIVQ spatial scores are significant predictors of classification accuracy. The finding that the OSIVQ spatial score constitutes a significant predictor here indicates that for classification a cruder retinotopic organization of mental imagery might already be sufficient. Hence successful classification may not be sufficient to draw conclusions regarding the geometry of imagined objects. As before, number of voxels also constitutes a significant predictor of classification accuracy.

The autoencoder enables leveraging perceptual data to improve reconstructions of imagined letters and pretrain classifiers. This may eventually be utilized for the development of content-based BCI letter-speller systems. So far, fMRI-based BCI communication systems have mostly focused on coding schemes arbitrarily mapping brain activity in response to diverse mental imagery tasks (e.g. mental spatial navigation, mental calculation, mental drawing or inner speech), and hence originating from distinct neural substrates, onto letters of the alphabet (Birbaumer et al. 1999; Sorger et al. 2012). As such, current BCI speller systems do not offer a meaningful connection between the intended letter and the specific content of mental imagery. This is demanding for users as it requires them to memorize the mapping in addition to performing imagery tasks. Our results constitute a proof-of-concept that it may be possible to achieve a more natural, content-based, BCI speller system immediately decoding internally visualized letters from their associated brain activity.

In conclusion, our letter encoding, reconstruction, and classification results indicate that the topographic organization of mental imagery closely resembles that of perception. This lends support to the idea that mental imagery is quasi-perceptual not only in terms of its subjective experience but also in terms of its neural representation and constitutes an important first step towards the development of content-based letter-speller systems.

## Acknowledgements

We would like to thank Carmine Gnolo for helpful discussions on the reconstruction procedure. This project has received funding from the European Union’s Horizon 2020 Research and Innovation Programme under Grant Agreement No. 7202070 (HBP SGA1) as well as under ERC-2010-AdG grant (269853).

## Compliance with ethical standards

### Studies involving human participants

All procedures were conducted with approval from the local Ethical Committee of the Faculty of Psychology and Neuroscience at Maastricht University.

### Informed consent

All participants gave written informed consent and were paid for participation in this study.

## References

Abadi M, Barham P, Chen J, Chen Z, Davis A, Dean J, Devin M, Ghemawat S, Irving G, Isard M, Kudlur M, Levenberg J, Monga R, Moore S, Murray DG, Steiner B, Tucker P, Vasudevan V, Warden P, Wicke M, Yu Y, Zheng X, Brain G. 2016. TensorFlow: A System for Large-Scale Machine Learning TensorFlow: A system for large-scale machine learning. In: 12th USENIX Symposium on Operating Systems Design and Implementation (OSDI ’16). p. 265–284.

Albers AMM, Kok P, Toni I, Dijkerman HCC, de Lange FP, de Lange FP. 2013. Shared Representations for Working Memory and Mental Imagery in Early Visual Cortex, Current Biology.

Andersson JLRR, Skare S, Ashburner J. 2003. How to correct susceptibility distortions in spin-echo echo-planar images: application to diffusion tensor imaging. Neuroimage. 20:870–888.

Birbaumer N, Ghanayim N, Hinterberger T, Iversen I, Kotchoubey B, Kübler A, Perelmouter J, Taub E, Flor H. 1999. A spelling device for the paralysed. Nature. 398:297–298.

Blazhenkova O, Kozhevnikov M. 2009. The new object-spatial-verbal cognitive style model: Theory and measurement. Appl Cogn Psychol. 23:638–663.

Brogaard B, Gatzia DE. 2017. Unconscious Imagination and the Mental Imagery Debate. Front Psychol. 8:799.

Cichy RM, Heinzle J, Haynes J-D. 2012. Imagery and Perception Share Cortical Representations of Content and Location. Cereb Cortex. 22:372–380.

Dijkstra N, Bosch SE, van Gerven MAJ. 2017. Vividness of Visual Imagery Depends on the Neural Overlap with Perception in Visual Areas. J Neurosci. 37:1367–1373.

Dumoulin SO, Wandell BAA. 2008. Population receptive field estimates in human visual cortex. Neuroimage. 39:647–660.

Emmerling TC, Zimmermann J, Sorger B, Frost MA, Goebel R. 2016. Decoding the direction of imagined visual motion using 7T ultra-high field fMRI. Neuroimage. 125:61–73.

Freeman J, Simoncelli EP. 2011. Metamers of the ventral stream. Nat Neurosci. 14:1195–1201.

Ganis G, Thompson WL, Kosslyn SM. 2004. Brain areas underlying visual mental imagery and visual perception: an fMRI study. Cogn Brain Res. 20:226–241.

Goebel R, Esposito F, Formisano E. 2006. Analysis of functional image analysis contest (FIAC) data with brainvoyager QX: From single-subject to cortically aligned group general linear model analysis and self-organizing group independent component analysis. Hum Brain Mapp. 27:392–401.

Goebel R, Khorram-Sefat D, Muckli L. 1998. The constructive nature of vision: direct evidence from functional magnetic resonance imaging studies of apparent motion and motion imagery. Eur J.

Harrison S, Tong F. 2009. Decoding reveals the contents of visual working memory in early visual areas. Nature.

Hummer A, Ritter M, Tik M, Ledolter AA, Woletz M, Holder GE, Dumoulin SO, Schmidt-Erfurth U, Windischberger C. 2016. Eyetracker-based gaze correction for robust mapping of population receptive fields. Neuroimage. 142:211–224.

Ishai A, Ungerleider L, Haxby J. 2000. Distributed neural systems for the generation of visual images. Neuron.

Johnson MR, Johnson MK. 2014. Decoding individual natural scene representations during perception and imagery. Front Hum Neurosci. 8:59.

Kingma DP, Ba J. 2014. Adam: A Method for Stochastic Optimization.

Kosslyn S, Thompson W, Alpert N. 1997. Neural systems shared by visual imagery and visual perception: A positron emission tomography study. Neuroimage.

Kozhevnikov M, Kozhevnikov M, Yu CJ, Blazhenkova O. 2013. Creativity, visualization abilities, and visual cognitive style. Br J Educ Psychol. 83:196–209.

Kriegeskorte N, Goebel R. 2001. An efficient algorithm for topologically correct segmentation of the cortical sheet in anatomical MR volumes. Neuroimage. 14:329–346.

Lee S, Kravitz D, Baker C. 2012. Disentangling visual imagery and perception of real-world objects. Neuroimage.

Marques JJP, Kober T, Krueger G, Zwaag W van der, van der Zwaag W, Van de Moortele P-F, Gruetter R. 2010. MP2RAGE, a self bias-field corrected sequence for improved segmentation and T 1-mapping at high field. Neuroimage. 49:1271–1281.

Mechelli A, Price C, Friston K, Ishai A. 2004. Where bottom-up meets top-down: neuronal interactions during perception and imagery. Cereb cortex.

Miyawaki Y, Uchida H, Yamashita O, Sato M, Morito Y. 2008. Visual image reconstruction from human brain activity using a combination of multiscale local image decoders. Neuron.

Moeller S, Yacoub E, Olman C. 2010. Multiband multislice GE-EPI at 7 tesla, with 16-fold acceleration using partial parallel imaging with application to high spatial and temporal whole-brain fMRI. Magnetic.

Naselaris T, Olman CA, Stansbury DE, Ugurbil K, Gallant JL. 2015. A voxel-wise encoding model for early visual areas decodes mental images of remembered scenes. Neuroimage. 105:215–228.

O’Craven K, Kanwisher N. 2000. Mental imagery of faces and places activates corresponding stimulus-specific brain regions. J Cogn Neurosci.

Pearson J, Naselaris T, Holmes E. 2015. Mental imagery: functional mechanisms and clinical applications. Trends Cogn.

Peirce J. 2007. PsychoPy—psychophysics software in Python. J Neurosci Methods.

Podgorny P, Shepard RN. 1978. Functional representations common to visual perception and imagination. J Exp Psychol Hum Percept Perform. 4:21–35.

Reddy L, Tsuchiya N, Serre T. 2010. Reading the mind’s eye: decoding category information during mental imagery. Neuroimage.

Schoenmakers S, Barth M, Heskes T, Gerven M van. 2013. Linear reconstruction of perceived images from human brain activity. Neuroimage.

Senden M, Reithler J, Gijsen S, Goebel R. 2014. Evaluating Population Receptive Field Estimation Frameworks in Terms of Robustness and Reproducibility. PLoS One. 9:e114054.

Slotnick S, Thompson W, Kosslyn S. 2005. Visual mental imagery induces retinotopically organized activation of early visual areas. Cereb cortex.

Smith S, Jenkinson M, Woolrich M, Beckmann C. 2004. Advances in functional and structural MR image analysis and implementation as FSL. Neuroimage.

Sorger B, Reithler J, Dahmen B, Goebel R. 2012. Report A Real-Time fMRI-Based Spelling Device Immediately Enabling Robust Motor-Independent Communication. Curr Biol. 22:1333–1338.

Stokes M, Saraiva A, Rohenkohl G, Nobre A. 2011. Imagery for shapes activates position-invariant representations in human visual cortex. Neuroimage.

Stokes M, Thompson R, Cusack R. 2009. Top-down activation of shape-specific population codes in visual cortex during mental imagery. J.

Thirion B, Duchesnay E, Hubbard E, Dubois J, Poline J-B, Lebihan D, Dehaene S. 2006. Inverse retinotopy: inferring the visual content of images from brain activation patterns. Neuroimage. 33:1104–1116.

Thomas NJT. 1999. Are Theories of Imagery Theories of Imagination? An Active Perception Approach to Conscious Mental Content. Cogn Sci. 23:207–245.

Vincent P, Larochelle H, Bengio Y, Manzagol P-A. 2008. Extracting and composing robust features with denoising autoencoders. In: Proceedings of the 25th international conference on Machine learning - ICML ’08. New York, New York, USA: ACM Press. p. 1096–1103.

Wandell BA, Winawer J. 2015. Computational neuroimaging and population receptive fields. Trends Cogn Sci.

